# Mechanochemical Coupling and Bi-Phasic Force-Velocity Dependence in the Ultra-Fast Ring ATPase SpoIIIE

**DOI:** 10.1101/224022

**Authors:** Ninning Liu, Gheorghe Chistol, Yuanbo Cui, Carlos Bustamante

## Abstract

Multi-subunit ring-shaped ATPases are molecular motors that harness chemical free energy to perform vital mechanical tasks such as polypeptide translocation, DNA unwinding, and chromosome segregation. Previously we reported the intersubunit coordination and stepping behavior of the hexameric ring-shaped ATPase SpoIIIE (Liu et al., 2015). Here we use optical tweezers to characterize the motor’s mechanochemistry. Analysis of the motor response to external force at various nucleotide concentrations identifies phosphate release as the likely force-generating step. Analysis of SpoIIIE pausing indicates that pauses are off-pathway events. Characterization of SpoIIIE slipping behavior reveals that individual motor subunits engage DNA upon ATP binding. Furthermore, we find that SpoIIIE’s velocity exhibits an intriguing bi-phasic dependence on force. We hypothesize that this behavior is an adaptation of ultra-fast motors tasked with translocating DNA from which they must also remove DNA-bound protein roadblocks. Based on these results, we formulate a comprehensive mechanochemical model for SpoIIIE.

## Introduction

Ring-shaped ATPases perform tasks such as nucleic acid unwinding and chromosome segregation by orchestrating the operation of their individual subunits (Liu et al., 2014a). Each ATPase subunit cycles through a series of chemical transitions (ATP binding, hydrolysis, ADP and Pi release) and mechanical events (substrate binding, power-stroke, substrate release). The coupling between chemical and mechanical transitions determines how individual subunits operate while the coordination between subunits determines how the entire ring ATPase functions. Understanding the operating principles of these molecular machines requires a mechanistic model of ring ATPases at the level of individual subunits and the entire complex.

SpoIIIE is a homo-hexameric ring ATPase tasked with segregating the *B.subtilis* genome during sporulation (Shin et al., 2015). Among ring ATPases, SpoIIIE and its *E.coli* homologue FtsK stand out as the fastest-known nucleic acid translocases, pumping DNA at an astonishing 4000-7000 bp/s. (Lee et al., 2012; Ptacin et al., 2006). We have recently characterized SpoIIIE’s inter-subunit coordination and presented evidence for a two-subunit translocation-escort model where one subunit actively translocates DNA while its neighbor passively escorts DNA (Liu et al., 2015). However, the detailed mechano-chemical coupling underlying the operation of ultra-fast ATPases like SpoIIIE/FtsK remains largely unknown. Here we used optical tweezers to interrogate DNA translocation by SpoIIIE and dissect the mechanochemical cycle of each subunit.

## Results

### SpoIIIE Generates Up to 50 pN of Mechanical Force

Experiments were conducted on an instrument consisting of an optical trap and a micropipette as described previously (Liu et al., 2015). Briefly, SpoIIIE and DNA were immobilized separately on polystyrene beads (Figure 1A) and brought into proximity, allowing SpoIIIE to engage DNA. In the presence of ATP, SpoIIIE translocated DNA, shortening the tether between the two beads. Experiments were performed either in passive mode – where the trap position is fixed (Figure 1B), or in constant-force mode – where DNA tension is held constant (Figure 1C). At saturating [ATP] and low opposing force (5 pN), SpoIIIE translocated DNA at ~4 kbp/s (Figure 1C), in agreement with previous studies (Liu et al., 2015; Ptacin et al., 2008). Translocation rates measured in passive mode were in excellent agreement with those measured in constant-force mode (Figure 1D). We find that SpoIIIE can operate against forces up to 50 pN (Figure 1B), similar to other dsDNA translocases, including FtsK and the DNA packaging motors from bacteriophages T4, λ, and φ29 (Fuller et al., 2007a; 2007b; Saleh et al., 2004; Smith et al., 2001).

**Figure 1:**
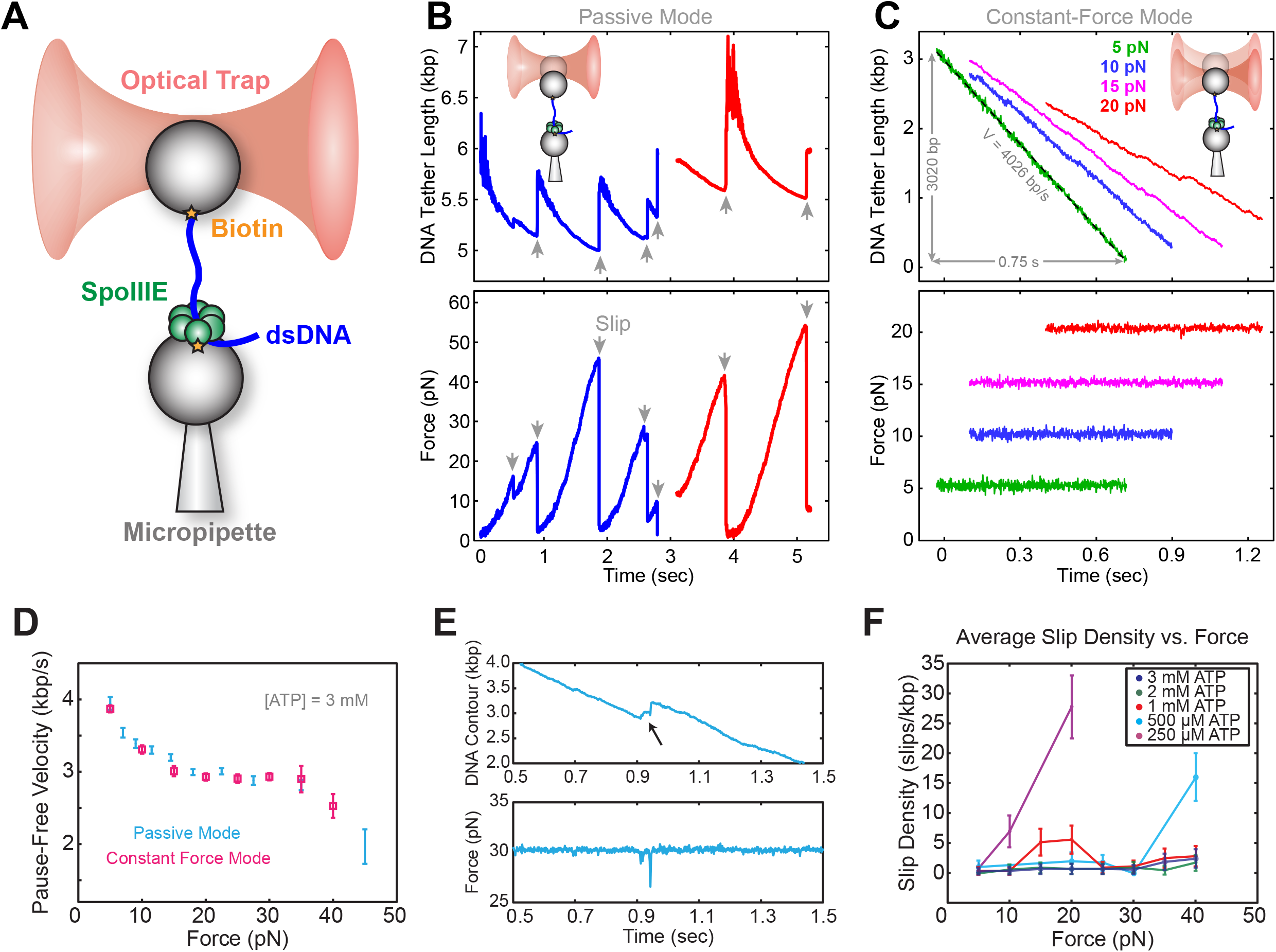
Optical tweezer experimental geometry in constant force and passive mode. (a) Optical tweezer geometry. (b) Representative single-molecule traces of SpoIIIE translocation in passive mode. The trap position is fixed and as SpoIIIE pulls the bead out of the trap the force on the trapped bead increases. (c) Representative single-molecule traces of SpoIIIE translocation in constant force mode. The optical trap position is continuously adjusted to maintain a constant force on the trapped bead. (d) Comparison of pause-free velocity measured in constant force mode and passive mode at [ATP] = 3 mM. Error-bars represent the standard error of the mean (SEM). (e) Trace displaying a slip in constant force mode. (f) Slip density at different opposing force and [ATP]. Error bars represent the square root of the number of events.

### ATP Mitigates Force-Induced Slipping

To investigate SpoIIIE operation, we monitored translocation in passive mode. At sufficiently high opposing forces (20-40 pN), translocation trajectories are often interrupted by slips, presumably due to SpoIIIE losing grip of its DNA track (Figure 1E). Eventually, SpoIIIE can recover, re-engage the DNA, and resume translocation from a low force; consequently, the same motor can undergo many rounds of continuous translocation and slipping.

As is shown in Figure 1-figure supplement 1A, at saturating [ATP] (3mM), SpoIIIE can undergo multiple rounds of pulling and slipping in passive mode, with a median slipping force of ~20pN (Figure 1-figure supplement 1B). At low [ATP], the median slipping force drops below 15 pN (Figure 1-figure supplement 1C), suggesting that the nucleotide state modulates the strength of SpoIIIE-DNA interactions. To investigate how [ATP] affects slipping, we conducted constant force experiments at 5-40 pN. At low [ATP] (0.25-0.50 mM) the slipping density increases sharply with opposing force, whereas at near-saturating [ATP] (1-3 mM) the slipping density is only weakly dependent on force (Figure 1F). Thus, binding of nucleotide to the motor appears to stabilize its grip on the DNA template.

### The SpoIIIE Power Stroke is Most Likely Driven by Pi Release

To probe how nucleotide binding is coordinated among motor subunits, we measured the pause-free SpoIIIE velocity at 3-50 pN of opposing force and 0.25-5.00 mM [ATP] (Figure 2A) using passive mode (Figure 2-figure supplement 1A). Fitting pause-free velocity versus [ATP] to the Hill equation, yields n_Hill_ = 1.0 ± 0.5 (Figure 2-figure supplement 1B-C). There are two means to achieve n_Hill_ ≈ 1 for a multi-subunit ATPase: (i) subunits turnover ATP independently of each other in an uncoordinated fashion; or (ii) subunits turnover ATP sequentially, but consecutive binding events are separated by an irreversible transition so only one subunit can bind nucleotide at any time, resulting in an *apparent* lack of cooperativity (Chemla et al., 2005). We recently found that SpoIIIE pauses when two neighboring subunits each bind a non-hydrolyzable ATP analog (Liu et al., 2015). This result is inconsistent with scenario (i) outlined above because an uncoordinated mechanism should enable several subunits to bind ATP analogs while the remaining subunits continue translocating. We conclude that SpoIIIE subunits bind ATP sequentially one subunit at a time. This coordination scheme enforces the well-defined subunit firing order required for SpoIIIE to track the backbone of one DNA strand as we previously showed (Liu et al., 2015).

**Figure 2:**
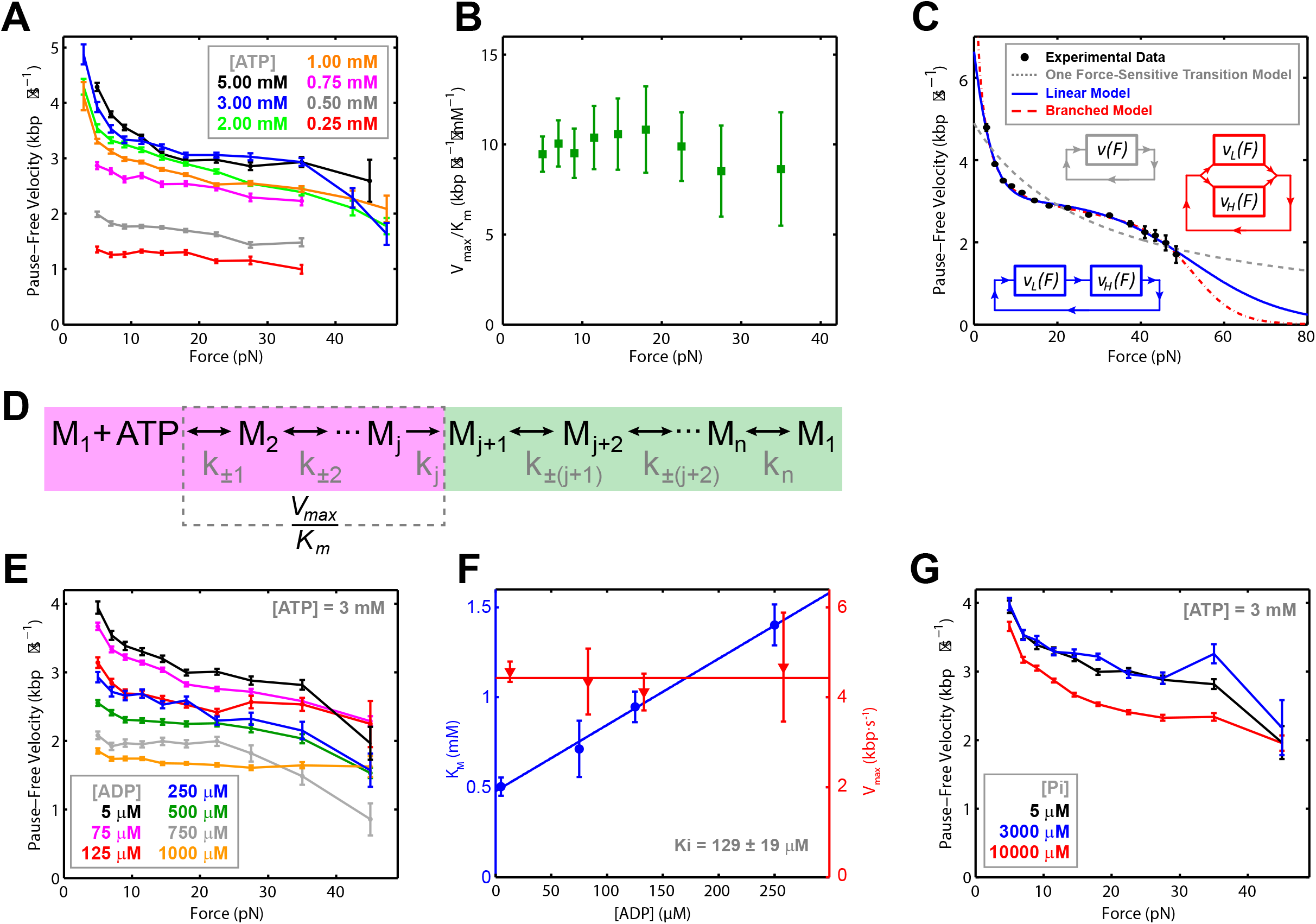
Force-velocity dependence displayed of SpoIIIE. (a) Pause-free translocation velocity versus opposing force at various [ATP], 5μM ADP, and 5μM P_i_. Error-bars represent the SEM. (b) Hill coefficient derived from fitting translocation velocity versus [ATP] at various opposing forces. Error bars represent the standard error of the fit (SEF). (c) Pause-free velocity versus opposing force (from data at 5, 3, and 2 mM ATP). Error-bars represent the SEM. Gray, blue and red curves represent fits to different models. Analytic expressions and fit parameters for the branched and linear models are given in panels Figure 2-figure supplement 2. (d) Generalized kinetic cycle for an ATPase subunit. The first block consists of all rate constants k_±1_, k_±2_,… up to the first irreversible transition kj (purple). The second block comprises the remaining rate constants (green). (e) Pause-free velocity versus opposing force at various [ADP] and 3 mM ATP. (f) V_max_ and K_M_ values as a function of [ADP] at low opposing force (5 pN). Solid lines are fits to a competitive inhibition model, K_i_ = 129 ± 19 μM. Error-bars represent the SEF. (g) Pause-free velocity versus opposing force under high [Pi] conditions and 3 mM ATP. Error bars represent the SEM.

To determine which chemical transition is coupled to the power stroke we investigated how force affects V_max_ and K_M_ determined from Michaelis-Menten fits. Although both V_max_ and K_M_ decrease with force, V_max_/K_M_ is largely force-independent (Figure 2B). To understand this result, consider a generalized ATPase cycle consisting of two kinetic blocks separated by an irreversible transition k_j_ (Figure 2D). We hypothesize that ATP tight binding (the transition that commits the ATPase to perform hydrolysis) is the irreversible transition that separates the kinetic blocks in Figure 2D, as has been proposed for other ring ATPases (Chemla et al., 2005; Moffitt et al., 2009; Sen et al., 2013). V_max_/K_M_ depends on ATP docking/undocking rates (k_±1_) and the rates of all kinetic transitions reversibly connected to ATP docking (k_±2_, k_±3_…) up to the first irreversible transition kj, (Figure 2D, purple) (Keller and Bustamante, 2000). The observed force-independence of V_max_/K_M_ indicates that ATP docking or any transition reversibly connected to it (Figure 2D, purple) cannot be the force-generating transition (Keller and Bustamante, 2000). Our observation that SpoIIIE is less force-sensitive at low [ATP], where nucleotide binding is rate-limiting, also suggests that ATP binding is not coupled to the power stroke, for if it were, the motor would be more, no less force sensitive, in rate-limiting conditions. Therefore, the force-generating transition must occur in the second block of the generalized kinetic cycle (Figure 2D, green). It is unlikely that ATP hydrolysis drives the power stroke because the rotation of the γ-phosphate upon hydrolysis does not release sufficient free energy (Oster and Wang, 2000). Therefore, ADP or P_i_ release — both of which are located in the second kinetic block (Figure 2D, green) — must be responsible for force generation.

To distinguish between these possibilities, we quantified the inhibitory effect of ADP and Pi on translocation. As [ADP] was titrated from 5 to 1000 μM the pause-free velocity decreased (Figure 2E, Figure 2-figure supplement 1D). The apparent K_M_ increases linearly with [ADP] whereas V_max_ is independent of [ADP] (Figure 2F, Figure 2-figure supplement 1E), indicating that ADP is a competitive inhibitor to ATP binding with a dissociation constant K_d_ = 129 ± 19 μM. In contrast, at the highest [P_i_] sampled (10 mM), the pause-free velocity decreased by only ~12% relative to the lowest [P_i_] tested (5 μM) (Figure 2G) indicating that P_i_ release is largely irreversible with a K_d_ >>10 mM. Given these Kd values, we estimated the change in free energy upon Pi and ADP release ΔG_Pi_ > 7.6 k_B_T and ΔG_ADP_ ~ 3.2 k_B_T in a buffer containing 5 μM P_i_ and 5 μM ADP (Chemla et al., 2005) (see Methods). For an estimated step size of 2 bp (Liu et al., 2015), and a maximum generated force of ~50 pN, each SpoIIIE power-stroke requires at least 8.2 k_B_T of free energy (see Methods). We conclude that P_i_ release is the only chemical transition capable of driving the power stroke of SpoIIIE, similar to what has been proposed for the φ29 packaging motor (Chemla et al., 2005), and the ClpX ring ATPase (Sen et al., 2013).

### The SpoIIIE Cycle Contains at Least Two Force-Dependent Kinetic Rates

At near-saturating [ATP], SpoIIIE exhibits a bi-phasic force-velocity dependence: the pause-free velocity drops rapidly between 5 and 15 pN, remains relatively force-insensitive between 15 and 40 pN, then rapidly decreases again beyond 40 pN (Figure 2A). Because force-induced slipping limited the amount of data above 40 pN, (Table 1), we combined the data at near-saturating [ATP] (2, 3, 5mM) into a consolidated curve (Figure 2C) that clearly displays the bi-phasic force-velocity dependence. A model with a single force-sensitive transition is inconsistent with this observation: (i) it predicts a monotonic decrease in velocity with force and poorly fits the data (Figure 2C, dashed gray curve), and (ii) it requires more free energy per power stroke than is released by hydrolyzing one ATP (see Methods).

**Table 1:** Length of DNA (kbp) translocated at different forces and ATP concentrations. Related to Figure 2C

| **[ATP]**<br>**(μM)** | ***Force Interval (pN)*** |  |  |  |  |  |  |  |  |  |  |  |
| --- | --- | --- | --- | --- | --- | --- | --- | --- | --- | --- | --- | --- |
|  | **2-4** | **4-6** | **6-8** | **8-10** | **10-13** | **13-16** | **16-20** | **20-25** | **25-30** | **30-40** | **40-45** | **45-50** |
| 5000 | 105.0 | 98.2 | 75.1 | 61.5 | 77.6 | 60.3 | 51.4 | 34.8 | 16.2 | 11.8 | 1.0 | 0.2 |
| 3000 | 44.6 | 40.1 | 33.7 | 30.0 | 42.4 | 38.6 | 45.5 | 42.6 | 23.7 | 16.3 | 1.9 | 0.6 |
| 2000 | 39.0 | 38.9 | 34.6 | 32.1 | 45.4 | 41.5 | 47.9 | 41.3 | 21.0 | 12.1 | 1.8 | 0.9 |
| 1000 | 76.5 | 66.3 | 58.6 | 54.9 | 77.1 | 70.1 | 76.9 | 60.2 | 29.6 | 19.1 | 1.6 | 0.5 |
| 750 | 23.8 | 18.4 | 14.4 | 12.3 | 16.6 | 14.1 | 14.1 | 10.1 | 5.2 | 3.6 | 0.3 | 0 |
| 500 | 44.1 | 43.7 | 37.8 | 33.3 | 42.4 | 34.9 | 31.5 | 19.6 | 8.1 | 3.7 | 0 | 0 |
| 250 | 22.4 | 20.4 | 18.4 | 17.4 | 23.4 | 17.1 | 13.5 | 9.9 | 4.2 | 2.0 | 0 | 0 |

At least two force-sensitive transitions are needed to rationalize SpoIIIE’s force-velocity dependence: the first should capture the motor’s sensitivity to force at low loads (<15 pN), the second should describe the motor’s sensitivity to force at high loads (>40 pN). Two kinetic schemes – branched and linear – are consistent with the SpoIIIE data. The branched scheme contains two parallel routes: at low forces the motor operates through the branch with high sensitivity to force, while at high forces it functions through the branch with low sensitivity to force (Figure 2-figure supplement 2). In contrast, the linear scheme contains two load-dependent transitions arranged in series (Figure 2-figure supplement 2). Although both kinetic schemes can fit the bi-phasic force-velocity dependence, the branched and linear schemes have fundamentally distinct implications for SpoIIIE operation. Based on studies of related ring ATPases, we favor the linear scheme (see Discussion).

### Pausing is in Kinetic Competition with Translocation and ATP Binding

At low [ATP], SpoIIIE exhibits spontaneous pausing (Figure 3A). We find that pause density increases dramatically as pause-free velocity drops (Figure 3B), suggesting that pausing and translocation are in kinetic competition. This observation is consistent with a model where the pause state is off the main translocation pathway. To determine where in the cycle the off-pathway pause state is located, we analyzed SpoIIIE’s pausing at various [ATP] and forces. We find that pause density increases drastically at low [ATP] (Figure 3C) indicating that pausing is in kinetic competition with nucleotide binding. In other words, SpoIIIE enters a pause when a subunit is awaiting ATP binding. We also found that the mean pause duration is inversely proportional to [ATP] (Figure 3D) suggesting that SpoIIIE exits the paused state by binding nucleotide. Pause durations at a given [ATP] are exponentially distributed (Figure 3D, inset) indicating that pause duration is governed by a single-rate limiting event—presumably the motor binding an ATP molecule. Finally, the fact that the pause density and pause duration does not depend on force at low [ATP] (where ATP binding is rate-limiting) (Figure 3E-F) suggests that the pause state is not reversibly connected to the force-generating transition (Pi release).

**Figure 3:**
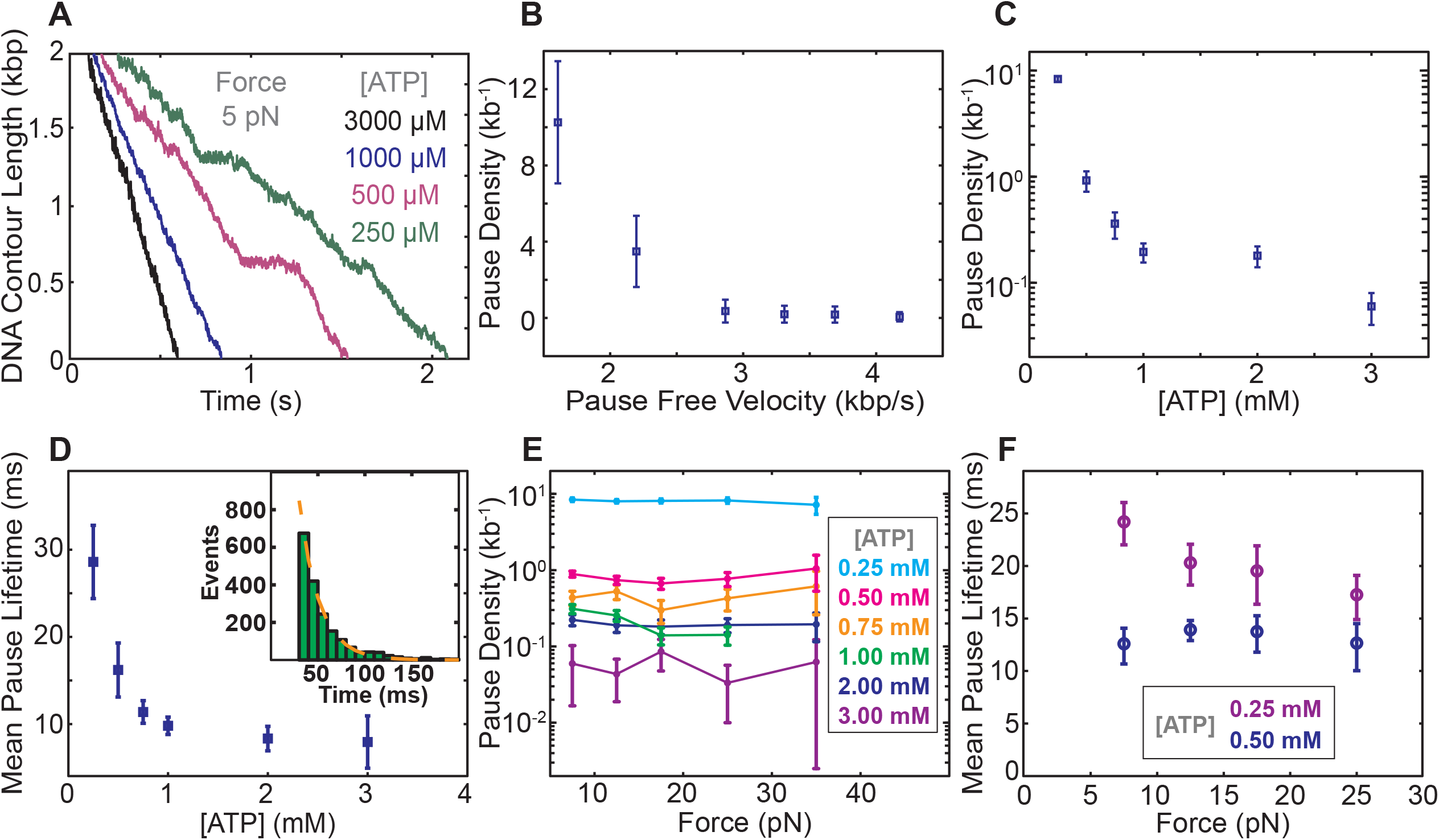
Characterization of spontaneous pausing by SpoIIIE (a) Examples of SpoIIIE pausing at low force (5 pN) and various ATP concentrations. (b) Pause density versus pause-free velocity at 5 pN of force. (c) Pause density versus ATP concentration at 5 pN of force. (d) The mean pause lifetime calculated by fitting the distribution of pause durations to a single exponential (see inset). Error-bars represent the SEF. (Inset) Distribution of pause durations at 250 μM [ATP] (green) fit to a single-exponential decay (dashed red line). (we were unable to accurately estimate the mean pause lifetime at high [ATP] due to the low number of detectable pauses) (e) Pause density versus opposing force at various [ATP]. (f) Mean pause lifetimes versus opposing force at the two lowest [ATP], where the number of pauses was sufficiently high to accurately estimate the lifetimes from fits. Error bars from fits represent 95% CI from fits. Error bars of pause density estimated from square root of the number of pause events.

## Discussion

### The Mechanochemical Cycle of an Individual SpoIIIE Subunit

Based on the results above, we propose a minimal mechanochemical model for a single SpoIIIE subunit (Figure 4A). The ADP-bound state (gray) is reversibly connected to the Apo state (white), which is reversibly connected to the ATP-loosely-docked state (light green). Here ADP acts as a competitive inhibitor to ATP binding, as observed experimentally. The ATP-loosely-docked state is irreversibly connected to the ATP-tightly-bound state (dark green), ensuring that V_max_/K_M_ is force-insensitive. Hydrolysis is depicted as a reversible process between the ATP-tightly-bound state and the transition state ADP · P_i_ (blue). Finally, P_i_ release is depicted as an irreversible process that drives the 2-bp power-stroke.

**Figure 4:**
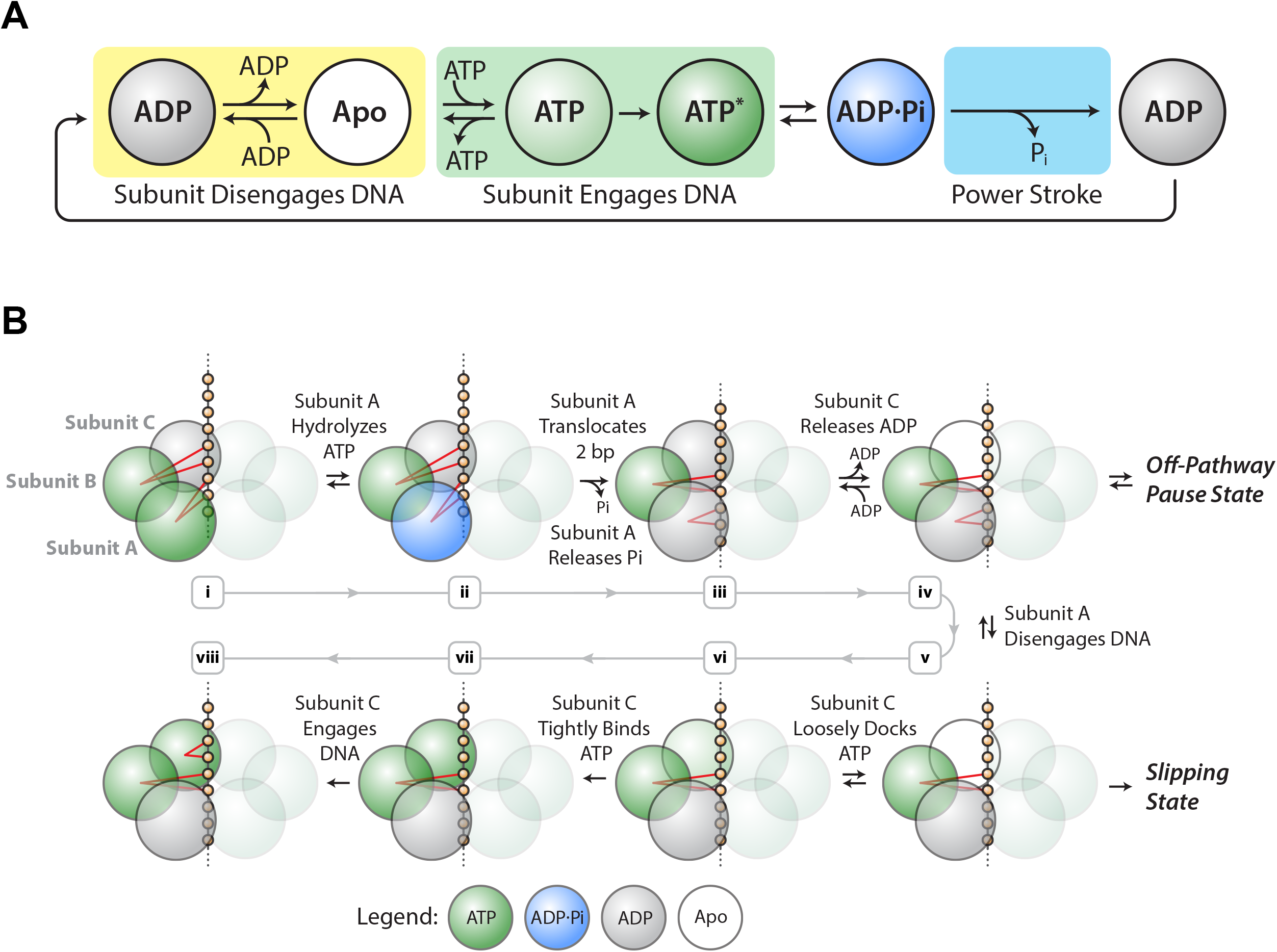
SpoIIIE mechanochemistry model (a) Mechanochemical cycle for a single SpoIIIE subunit. (b) Mechanochemical cycle for the entire SpoIIIE homo-hexamer.

When does SpoIIIE make and break its DNA contacts? We previously found that the strength of the motor-DNA interaction is highest in the ATPγS-bound state, moderate in the ADP-bound state, and lowest in the Apo state (Liu et al., 2015). Given that nucleotides strengthen the motor-DNA interactions, we propose that each SpoIIIE subunit has to bind ATP first before it engages the DNA and the motor-DNA interaction is established during ATP docking or during tight-binding (Figure 4A, green box). Since the ADP-bound and the Apo states have the weakest affinity for DNA we propose that each subunit breaks its contacts with DNA after reaching the ADP-bound state or the Apo state (Figure 4A, yellow box).

### Force-Induced Slipping: Implications for the Two-Subunit Translocation-Escort Mechanism

We previously provided evidence for a model where two subunits contact the DNA at adjacent pairs of phosphates on the same strand: while one subunit executes the power-stroke and translocates 2 bp, the other escorts the DNA (Liu et al., 2015). This mechanism enables the motor to operate processively with non-consecutive inactive subunits, and the escorting subunit may function as a backup should the translocating subunit loose its grip on DNA during the power-stroke. In the present study, we find that (1) force-induced slipping is more frequent at low [ATP], and (2) at low [ATP] the slipping probability increases under higher opposing force.

Based on the insights from the slipping data, we propose a revised model of the one proposed before (Liu et al., 2015) (Figure 4B). After executing the power-stroke, the translocating subunit (A) disengages DNA, the escorting subunit (B) maintains its grip on DNA while the next subunit (C) first binds ATP and then engages the DNA (Figure 4Biii-vi). After this hand-over, subunits B and C become the new translocating and escorting subunits respectively and this cycle continues around the ring. The motor is most vulnerable to slipping while B is the only subunit anchoring the hexamer to the DNA backbone (Figure 4Bv). At high [ADP] and low [ATP], subunit C spends more time in the ADP-bound or Apo state, lengthening the time in which subunit B is the only one anchoring the motor onto DNA, and increasing the slipping probability (Figure 2-figure supplement 3). When the escorting and translocating subunits are both contacting DNA, the likelihood of force-induced slipping is significantly diminished.

In our model ADP release happens before ATP binding (see subunit C in Figure 4Biii-iv). Since ADP acts as a competitive inhibitor to ATP binding, in our model ADP release and ATP docking in the same subunit are connected via reversible transitions as depicted in Figure 4Biii-vi. It is unclear what triggers ADP release, however studies of related ATPases show that ADP release is highly coordinated among subunits, triggered for example by the binding of ATP in the adjacent subunit (Chistol et al., 2012).

### Off-Pathway Pausing: Timing and Implications

We found that pause density is inversely proportional to pause-free velocity (Figure 3B), indicating that pausing is an off-pathway process in kinetic competition with translocation. The observation that pausing is more likely at low [ATP] (Figure 3E) suggests that SpoIIIE pauses when a subunit is awaiting the binding of ATP. At the same time, we do not observe frequent slipping from paused states. We speculate that SpoIIIE enters off-pathway pauses from the state depicted in Figure 4Biv – after subunit A translocated but before DNA is handed to subunit C. At this stage, subunit C is poised to bind ATP. At low [ATP] the motor may detect if subunit C takes a long time to bind nucleotide and the SpoIIIE would enter the off-pathway pause state while gripping the DNA with two subunits. Such an allosteric sensing mechanism would prevent the motor from prematurely initiating the slip-prone DNA handover (Figure 4Bv). This speculative regulatory mechanism for SpoIIIE is reminiscent of the allosteric regulation of the φ29 viral packaging motor, which senses when the capsid is nearly full and enters into long-lived pauses allowing DNA inside the capsid to relax before packaging can restart (Berndsen et al., 2015; Liu et al., 2014b). During chromosome segregation in *B. subtilis,* the local [ATP] near active SpoIIIE complexes could fluctuate on short time-scales. A drop in local [ATP] could force the motor to pause thus preventing the slip-prone handover until [ATP] rises to levels optimal for SpoIIIE operation.

### Bi-Phasic Velocity-vs-Force Dependence and Its Implications

In addition to SpoIIIE, bi-phasic force-velocity dependence has been reported for two other ultra-fast DNA translocases — FtsK (Saleh et al., 2004), and λ phage packaging motor (Fuller et al., 2007b). This unusual behavior can be explained by linear or branched models with two force-dependent transitions. A plausible branched model is one where the force-sensitive transitions represent two alternative power strokes, one predominant at low loads with another predominant at high loads, and the motor executes only one of these transitions in each cycle (Figure 2-figure supplement 2). Although there are molecular motors with two power strokes that are executed sequentially (Adachi et al., 2007), no motor has been shown to perform one at the exclusion of the other.

Thus, we favor the linear model where the force-dependence above 40 pN corresponds to the force-sensitivity of the power-stroke. We speculate that the force-dependence observed at low force reflects a load-induced motor deformation that slows down a kinetic transition distinct from the power stroke, and this deformation saturates at ~15 pN. The force-independence of V_max_/K_M_ suggests that the low-force, force-dependent transition occurs after ATP tight binding (ATP hydrolysis, ADP release, or another transition distinct from P_i_ release). Finally the linear model predicts a velocity of ~6.5 kbp/s under no load (Figure 2-figure supplement 2), in agreement with the zero-force maximum velocity of FtsK – SpoIIIE’s homologue in *E. coli* (Lee et al., 2012).

Interestingly, the *in vivo* SpoIIIE rate is markedly slower (~1-2 kb/sec) than its *in vitro* rate (~4-5 kb/sec) (Burton et al., 2007; Ptacin et al., 2008). This discrepancy can be explained by the fact that DNA-bound proteins act as physical barriers to SpoIIIE *in vivo* effectively creating an opposing force (Marquis et al., 2008). Although we do not have an accurate estimate of the opposing forces experienced by individual motors *in vivo*, we expect that they are up to tens of pN because such forces are needed to disrupt protein-DNA interactions *in vitro* (Dame et al., 2006). Our study provides insight into how ultrafast ring ATPases like SpoIIIE and FtsK may respond to a variety of physical and chemical challenges inside the cell, such as decreasing translocation velocity when encountering opposing forces, slipping at high forces, and pausing at low ATP concentrations.

### Materials and Methods

Sample Preparation

Recombinant SpoIIIE, dsDNA substrates, and polystyrene beads were prepared as described before (Liu et al., 2015).

### Data Analysis

Pauses were detected using a modified Schwartz Information Criterion (mSIC) method (Maillard et al., 2011). The number and duration of pauses missed by this algorithm were inferred by fitting the pause durations to a single exponential with a maximum likelihood estimator (Liu et al., 2015). After removing the detected pauses, the translocation velocity was computed by fitting the data to a straight line. For passive-mode data, singlemolecule trajectories were partitioned into segments spanning 2-3 pN, and the velocity was computed for each segment. Tether tension and extension were converted to contour length using the Worm-Like-Chain approximation (Baumann et al., 1997).

### Estimating Free Energy of Product Release

To estimate the free energy of product release, consider the simplified kinetic scheme *E · P ⇆ E + P* where the enzyme (E) can release or bind its product (P) with a forward and reverse rate krel and kbind respectively. We can define the rate of phosphate release as k_rel_ = k_-p_ and the rate of phosphate binding as k_bind_ = k_p_ [P_i_] where k_-p_ and k_p_ are the first and second-order rate constants for phosphate release and binding respectively. The free energy change corresponding to phosphate release is given by ΔG_Pi_ = – k_B_T·ln(k_rel_/k_bind_) = – k_B_T·ln(k_-p_/k_p_·[P_i_]) (Chemla et al., 2005). Since concentrations of phosphate as high as 10mM do not significantly affect SpoIIIE’s translocation velocity, then k_rel_ must be significantly higher than k_bind_ at P_i_ concentrations of 10 mM or less (i.e., k_-p_>> k_p_·[10 mM]) >>1. From these inequalities we can infer that k_-p_ /(k_p_[5 μM]) > 2000 and therefore we can set a lower bound for the free energy of P_i_ release as ΔG_Pi_ > – 7.6 k_B_T in a buffer containing 5 μM P_i_.

Similarly, we used the equilibrium dissociation constant for ADP (K_ADP_ = 129 ± 19 μM) to estimate the change in free energy associated with ADP release: ΔG_ADP_ ~ 3.2 k_B_T in standard buffer conditions ([ADP] = 5 μM). Given the estimated SpoIIIE step size of 2 bp (Graham et al., 2010; Liu et al., 2015; Massey et al., 2006) and the fact that SpoIIIE can translocate DNA against forces as high as 50 pN each power-stroke requires a change in free energy of at least 50 pN · 2 bp · 0.34 nm/bp = 34 pN·nm = 8.2 k_B_T.

### Mechanochemical Model with a Single Force-Generating Transition

Such a model predicts that at saturating [ATP], the motor velocity (V) depends on the external load (*F*) as 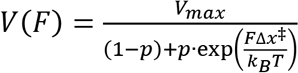 (Wang et al., 1998). Here *V_max_* is the maximum velocity at zero force, exp 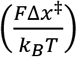 is an Arrhenius-like term describing how the external load slows down the force-generating transition, *p* is the fraction of the total mechanochemical cycle time that the motor spends in the force-generating transition at zero force, (*1-p*) captures all the force-independent transitions from the motor’s cycle, *k_BT_* is the Boltzmann constant times the temperature, and Δ*x*^‡^ is the distance to the transition state for the force-generating transition.

Fitting the consolidated force-velocity curve to the model above produces a very poor fit to the data (Figure 2C, dashed gray curve) (*V_max_* = 4.2 ± 0.4 kbp/s), Δ*x*^‡^ = 0.07 ± 0.02 nm, p ≈ 1), and most importantly predicts a monotonic decrease in velocity with force that does not capture the bi-phasic force-velocity dependence exhibited by SpoIIIE. Furthermore, extrapolating the fit to higher forces predicts large translocation velocities (>300 bp/sec) for loads over 400 pN. Considering that the likely step size of SpoIIIE is 2 bp per nucleotide hydrolyzed (Liu et al., 2015), a stall force above 400 pN requires that the motor generate at least 400 pN · 2 bp · 0.34 nm/bp = 270 pN·nm of work per power-stroke—more than two and a half times the ~110 pN · nm of free energy available from ATP hydrolysis in our experiments.

### Deriving Expressions for the Branched and Linear Models of Force-Velocity Dependence

We considered two broad classes of kinetic models that can capture the bi-phasic velocity dependence on force: branched models and linear models. In each case the average time needed to complete one cycle can be computed given the rate of ATP binding (which is proportional to [ATP] with the proportionality constant *α*) the rates of the two force-sensitive kinetic transitions (*k_A_* and *k_B_*), and the net compound rate of all remaining kinetic transitions that are force-insensitive (*k_0_*). The average cycle completion time for the branched model shown in Figure 2B-figure supplement 2A, *τ_branched_*, can be written as follows:

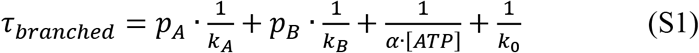

Here *p_A_* and *p_B_* are the probabilities that the cycle proceeds through each of the two force-sensitive branches (Figure 2B-figure supplement 2A). For simplicity we assumed an Arrhenius-like dependence on force *F* for *p_A_*.

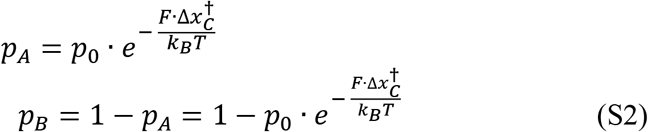

The rates for the two force-sensitive transitions are given by *k_A_* and *k_B_* respectively, each with an Arrhenius-like dependence on force *F* as shown below. Here *k_B_T* is the product of the Boltzmann constant and the temperature, *k_A0_* and *k_B0_* are the rates at zero force, and *Δx^†^_A_* and *Δx^†^_B_* are the distances to the transition state for the two force-sensitive branches.

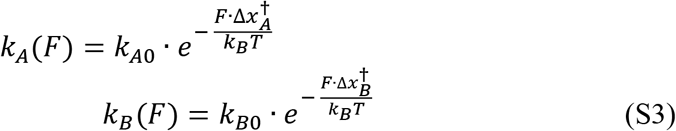

Each of the two force-sensitive transitions represents a power-stroke with step-sizes *d_A_* and *d_B_* respectively. Therefore, the average step size for the branched cycle *d_branched_* is given by:

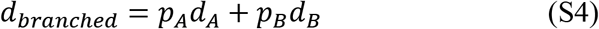

We fit the force-velocity data in Figure 2C to the simplest branched model where *d_A_ = d_B_* = *d_branched_ = d*. Note that the model in which *d_A_ Φ d_B_* also fits the data, but is less well-constrained. The average translocation velocity for the branched model is given by the following expression:

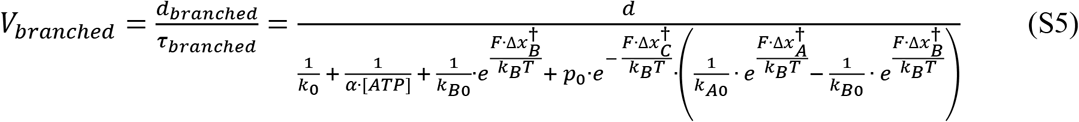

In a similar fashion, an expression for the translocation velocity can be derived for the linear model depicted in Figure 2C.

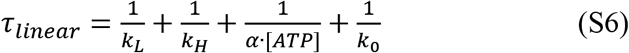

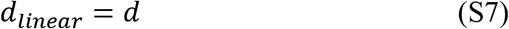

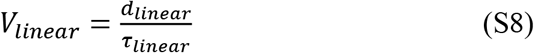

Here *kL* and *kH* are the rates of the force-sensitive transitions responsible for the drop in velocity at low forces (0-15 pN) and high forces (40 pN and above). *k_H_* represents the rate of the force-generating transition, i.e. phosphate release (most likely), and is given by a simple Arrhenius-like dependence:

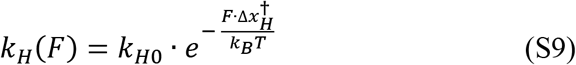

As described in the main text, *k_L_* saturates at a certain force (~15 pN), and could be written as follows:

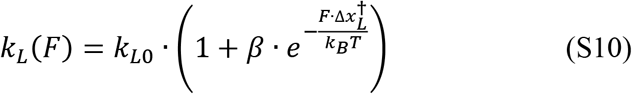

The final expression for *Vlinear* is:

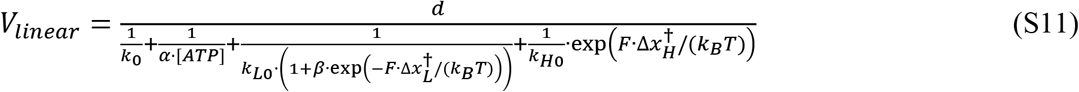

### Fitting the Consolidated Force-Velocity Curve to The Linear Model

The linear model provides two values for the distance to the transition state, 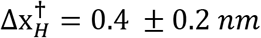 at high forces and 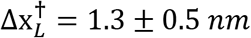 at low forces. A typical energy landscape for a molecular motor contains both a chemical axis, which captures the sequential chemical transitions a motor undergoes as it generates mechanical work, and a mechanical axis, which captures the physical movement of the motor along a distance coordinate (Bustamante et al., 2004). The distance to the transition state Δx^†^ is the distance the motor must move along the mechanical coordinate during the force-sensitive step in order to commit itself to stepping. If a motor directly couples a chemical transition to the force-generating step, in what is classically referred to as a “power stroke”, the motor will move approximately along a diagonal across the chemical and mechanical axes, and Δx_†_ would typically be < Δx_step_, where Δx_step_ is the distance the motor moves per step size. The value for 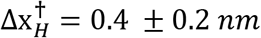 is smaller than and consistent with a 2-bp step size (0.68 nm) power stroke mechanism previously determined for SpoIIIE (Liu et al., 2015) and likely coupled P_i_ release. The other distance to the transition state, 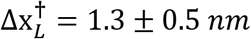, captures the hypothesized deformation of the protein at low forces.

## Acknowledgements

We thank S. B. Smith for instrument training and assistance; J. Y. Shin, M. Righini, B. Onoa, and C. Diaz for discussions. This work was supported by NIH grants R01GM032543 and the U.S. Department of Energy Office of Basic Energy Sciences Nanomachine Program under contract no. DE-AC02-05CH11231.

## Author Contributions

N.L., G.C., and C.B. conceived the project and designed the experiments; N.L. conducted the majority of single-molecule experiments; Y.C. collected data in constant-force mode; N.L. and G.C. prepared samples and analyzed the data; G.C. and N.L. wrote MATLAB code for data analysis; N.L., G.C., Y.C. and C.B. wrote the manuscript.

---

**Figure 1-figure supplement 1:**
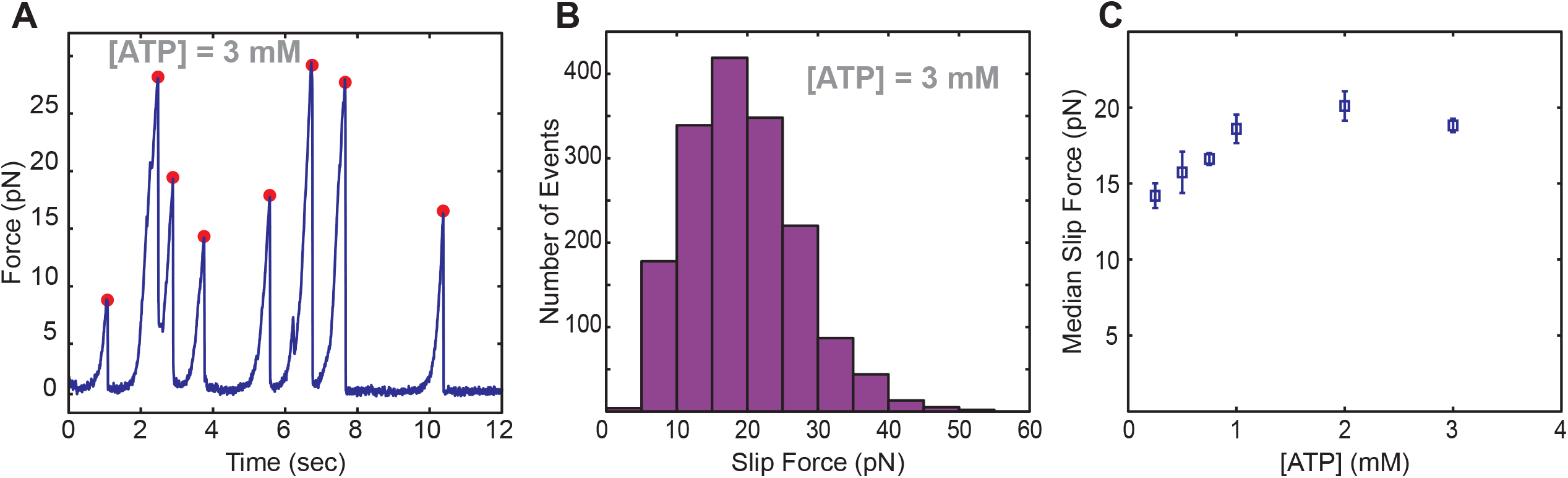
Slipping behavior of SpoIIIE. (a) Single-molecule trace of SpoIIIE translocation acquired in passive mode. Red dots indicate force where a slip was detected. (b) Histogram of pull forces. (c) Median pull force of SpoIIIE across various ATP conditions. Error bars display the standard error estimated from bootstrapping.

**Figure 2-figure supplement 1.**
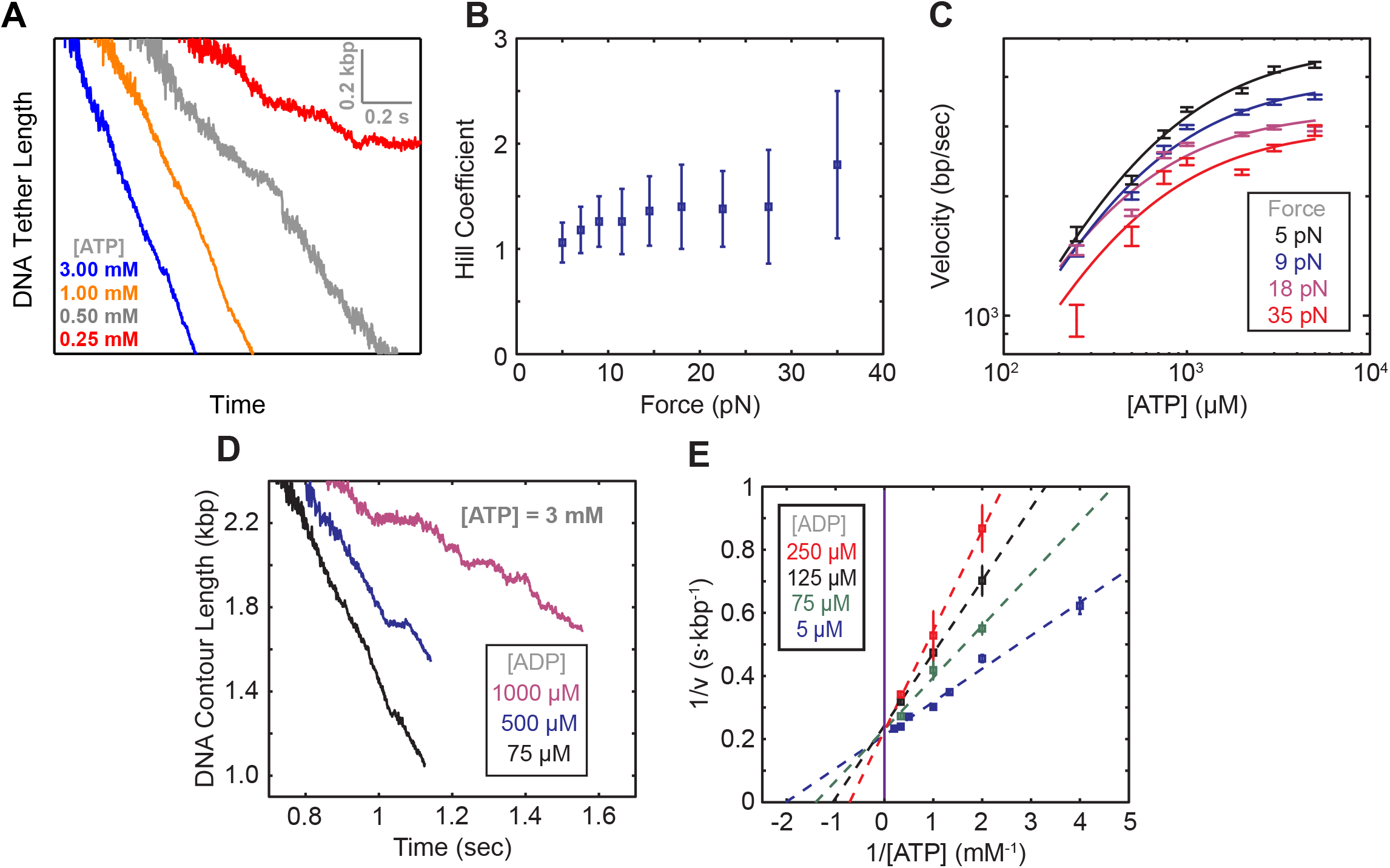
(a) Passive mode traces of SpoIIIE translocation across various [ATP]. (b) Hill coefficient derived from fitting translocation velocity versus [ATP] at various opposing forces. Error bars represent the standard error of the fit (SEF). (c) Examples of Michaelis-Menten fits to translocation data at different opposing forces. (d) Representative translocation traces acquired in passive mode at 3 mM ATP and various ADP concentrations. Error-bars represent the SEM. (e) Lineweaver-Burke plots at various [ADP] (v denotes the pause-free velocity). Dotted lines represent the Michaelis-Menten fits. The solid purple line marks the y-intercept. Error bars represent the SEM.

**Figure 2-figure supplement 2.**
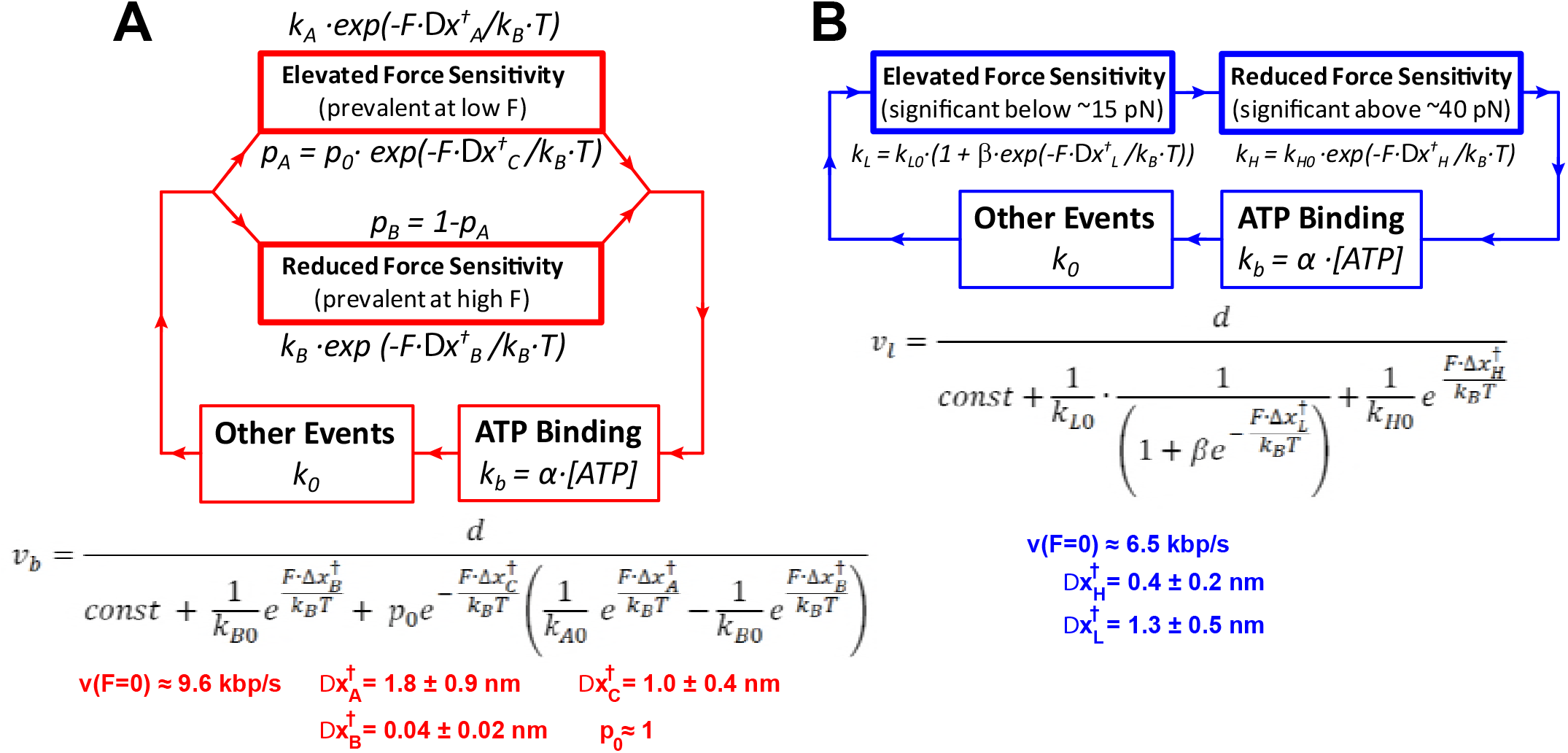
(a) Diagram illustrating the branched model. (b) Diagram illustrating the linear model.

**Figure 2-figure supplement 3.**
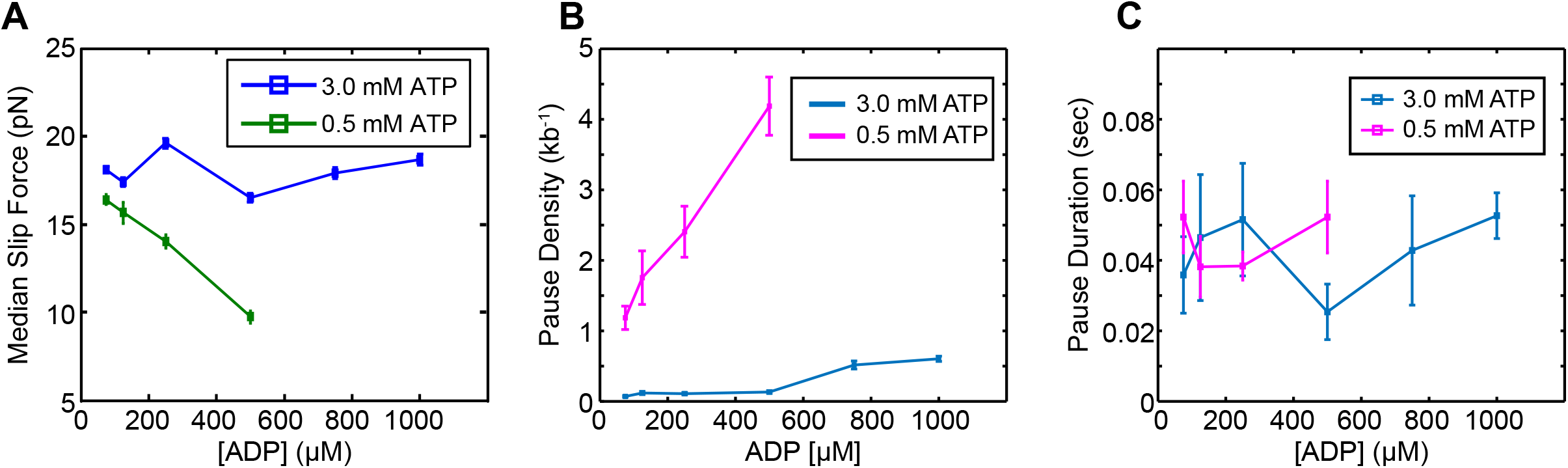
(a) Median slip force of SpoIIIE as a function of [ADP] at high (3 mM) and low (0.5 mM) [ATP]. Error bars represent the SEM. (b) Pause density versus [ADP] at both high and low ATP concentrations. Error bars display the square root of the pause number. (c) Mean pause durations calculated from single-exponential fits versus ADP concentration at high and low [ATP]. Error bars represent the 95% CI of the fit.

## References

Adachi, K., Oiwa, K., Nishizaka, T., Furuike, S., Noji, H., Itoh, H., Yoshida, M., and Kinosita, K. (2007). Coupling of rotation and catalysis in F(1)-ATPase revealed by single-molecule imaging and manipulation. Cell 130, 309–321.

Baumann, C.G., Smith, S.B., Bloomfield, V.A., and Bustamante, C. (1997). Ionic effects on the elasticity of single DNA molecules. Proceedings of the National Academy of Sciences 94, 6185–6190.

Berndsen, Z.T., Keller, N., and Smith, D.E. (2015). Continuous allosteric regulation of a viral packaging motor by a sensor that detects the density and conformation of packaged DNA. 108, 315–324.

Burton, B.M., Marquis, K.A., Sullivan, N.L., Rapoport, T.A., and Rudner, D.Z. (2007). The ATPase SpoIIIE transports DNA across fused septal membranes during sporulation in Bacillus subtilis. Cell 131, 1301–1312.

Bustamante, C., Chemla, Y.R., Forde, N.R., and Izhaky, D. (2004). Mechanical processes in biochemistry. Annu. Rev. Biochem. 73, 705–748.

Chemla, Y.R., Aathavan, K., Michaelis, J., Grimes, S., Jardine, P.J., Anderson, D.L., and Bustamante, C. (2005). Mechanism of Force Generation of a Viral DNA Packaging Motor. Cell 122, 683–692.

Chistol, G., Liu, S., Hetherington, C.L., Moffitt, J.R., Grimes, S., Jardine, P.J., and Bustamante, C. (2012). High degree of coordination and division of labor among subunits in a homomeric ring ATPase. Cell 151, 1017–1028.

Dame, R.T., Noom, M.C., and Wuite, G.J.L. (2006). Bacterial chromatin organization by H-NS protein unravelled using dual DNA manipulation. Nature 444, 387–390.

Fuller, D.N., Raymer, D.M., Kottadiel, V.I., Rao, V.B., and Smith, D.E. (2007a). Single phage T4 DNA packaging motors exhibit large force generation, high velocity, and dynamic variability. Proceedings of the National Academy of Sciences 104, 16868–16873.

Fuller, D.N., Raymer, D.M., Rickgauer, J.P., Robertson, R.M., Catalano, C.E., Anderson, D.L., Grimes, S., and Smith, D.E. (2007b). Measurements of single DNA molecule packaging dynamics in bacteriophage lambda reveal high forces, high motor processivity, and capsid transformations. Journal of Molecular Biology 373, 1113–1122.

Graham, J.E., Sherratt, D.J., and Szczelkun, M.D. (2010). Sequence-specific assembly of FtsK hexamers establishes directional translocation on DNA. Proc. Natl. Acad. Sci. U.S.a. 107, 20263–20268.

Keller, D., and Bustamante, C. (2000). The mechanochemistry of molecular motors.

Lee, J.Y., Finkelstein, I.J., Crozat, E., Sherratt, D.J., and Greene, E.C. (2012). Single-molecule imaging of DNA curtains reveals mechanisms of KOPS sequence targeting by the DNA translocase FtsK. Proc. Natl. Acad. Sci. U.S.a. 109, 6531–6536.

Liu, N., Chistol, G., and Bustamante, C. (2015). Two-subunit DNA escort mechanism and inactive subunit bypass in an ultra-fast ring ATPase. Elife 4.

Liu, S., Chistol, G., and Bustamante, C. (2014a). Mechanical operation and intersubunit coordination of ring-shaped molecular motors: insights from single-molecule studies. 106, 1844–1858.

Liu, S., Chistol, G., Hetherington, C.L., Tafoya, S., Aathavan, K., Schnitzbauer, J., Grimes, S., Jardine, P.J., and Bustamante, C. (2014b). A viral packaging motor varies its DNA rotation and step size to preserve subunit coordination as the capsid fills. Cell 157, 702–713.

Maillard, R.A., Chistol, G., Sen, M., Righini, M., Tan, J., Kaiser, C.M., Hodges, C., Martin, A., and Bustamante, C. (2011). ClpX(P) generates mechanical force to unfold and translocate its protein substrates. Cell 145, 459–469.

Marquis, K.A., Burton, B.M., Nöllmann, M., Ptacin, J.L., Bustamante, C., Ben-Yehuda, S., and Rudner, D.Z. (2008). SpoIIIE strips proteins off the DNA during chromosome translocation. Genes & Development 22, 1786–1795.

Massey, T.H., Mercogliano, C.P., Yates, J., Sherratt, D.J., and Löwe, J. (2006). Double-stranded DNA translocation: structure and mechanism of hexameric FtsK. Molecular Cell 23, 457–469.

Moffitt, J.R., Chemla, Y.R., Aathavan, K., Grimes, S., Jardine, P.J., Anderson, D.L., and Bustamante, C. (2009). Intersubunit coordination in a homomeric ring ATPase. Nature 457, 446–450.

Oster, G., and Wang, H. (2000). Reverse engineering a protein: the mechanochemistry of ATP synthase. Biochim. Biophys. Acta 1458, 482–510.

Ptacin, J.L., Nöllmann, M., Becker, E.C., Cozzarelli, N.R., Pogliano, K., and Bustamante, C. (2008). Sequence-directed DNA export guides chromosome translocation during sporulation in Bacillus subtilis. Nature Publishing Group 15, 485–493.

Ptacin, J.L., Nöllmann, M., Bustamante, C., and Cozzarelli, N.R. (2006). Identification of the FtsK sequence-recognition domain. Nat Struct Mol Biol 13, 1023–1025.

Saleh, O.A., Pérals, C., Barre, F.-X., and Allemand, J.-F. (2004). Fast, DNA-sequence independent translocation by FtsK in a single-molecule experiment. The EMBO Journal 23, 2430–2439.

Sen, M., Maillard, R.A., Nyquist, K., Rodriguez-Aliaga, P., Pressé, S., Martin, A., and Bustamante, C. (2013). The ClpXP protease unfolds substrates using a constant rate of pulling but different gears. Cell 155, 636–646.

Shin, J.-Y., Lopez-Garrido, J., Lee, S.-H., Diaz-Celis, C., Fleming, T., Bustamante, C., and Pogliano, K. (2015). Visualization and functional dissection of coaxial paired SpoIIIE channels across the sporulation septum. Elife 4, e06474.

Smith, D.E., Tans, S.J., Smith, S.B., Grimes, S., Anderson, D.L., and Bustamante, C. (2001). The bacteriophage straight phi29 portal motor can package DNA against a large internal force. Nature 413, 748–752.

Wang, M.D., Schnitzer, M.J., Yin, H., Landick, R., Gelles, J., and Block, S.M. (1998). Force and velocity measured for single molecules of RNA polymerase. Science 282, 902–907.

